# Estimating full-field displacement in biological images using deep learning

**DOI:** 10.1101/2024.05.21.595161

**Authors:** Solomon J. E. T. Warsop, Soraya Caixeiro, Marcus Bischoff, Jochen Kursawe, Graham D. Bruce, Philip Wijesinghe

## Abstract

The estimation of full-field displacement between biological image frames or in videos is important for quantitative analyses of motion, dynamics and biophysics. However, the often weak signals, poor biological contrast and a multitude of noise processes typical to microscopy make this a formidable challenge for many contemporary methods. Here, we present a deep-learning method, termed Displacement Estimation FOR Microscopy (DEFORM-Net), that outperforms traditional digital image correlation and optical flow methods, as well as recent learned approaches, offering simultaneous high accuracy, spatial sampling and speed. DEFORM-Net is experimentally unsupervised, relying on displacement simulation based on a random fractal Perlin-noise process and optimised training loss functions, without the need for experimental ground truth. We demonstrate its performance on real biological videos of beating neonatal mouse cardiomyocytes and pulsed contractions in *Drosophila* pupae, and in various microscopy modalities. We provide DEFORM-Net as open source, including inference in the ImageJ/FIJI platform, for rapid evaluation, which will empower new quantitative applications in biology and medicine.

## 1 Introduction

Biological tissues are highly dynamic and motion is critical to their form and function [1]. As such, imaging of live samples at high spatiotemporal resolutions and over long term is coming to the fore in revealing a host of dynamic biophysical processes [2]. This emerging domain of imaging, however, is accompanied by new challenges. Namely, the accurate estimation of motion from biological images or videos at the relevant spatial and temporal resolutions, which is required to quantify biophysics. The high throughput of image data, limits in signal-to-noise ratios, and the diversity of biological contrast makes deep learning an attractive prospect to tackle this challenge.

The broad problem of recovering displacement from biological images can be classed in two categories. First, methods that seek to track individual objects over the field of view, such as particle, cell or organism tracking [3]. These methods have received much attention because of the recent advances in super-resolution microscopy at high spatiotemporal scales. They can help reveal biophysics from protein kinetics to collective organism behaviour. Recent methods have utilised deep learning because of its natural affinity for feature identification tasks [3]. Second, methods that seek to estimate the full-field displacement (displacement vector field), i.e., map the local displacement vectors at every spatial location between image frames or videos [4]. In contrast to object tracking, full-field displacement estimation is a necessary step towards quantifying tissue mechanics from the principles of continuum mechanics, which can reveal local forces, deformation and mechanical properties [5]. The estimation of full-field displacement has been utilised for various bio-applications [6, 7, 8, 9, 10, 11, 12, 13]; however, when compared to object tracking, these methods have not been specialised for live-imaging microscopy data.

Accurate estimation of full-field displacement between two image frames is typically carried out using digital image correlation (DIC) [14]. DIC solves the problem of: what displacement field leads to the greatest cross-correlation coefficient between reference and deformed image intensities? This registration procedure assumes that the local image textures are preserved when deformed. Because the solution to this problem is not unique, cross-correlation is maximised iteratively over many small subsets or patches of the images, the deformation of which is typically constrained to some affine transform with enforcement of continuity in displacement [6]. DIC has been developed and utilised primarily for experimental mechanics, such as materials testing and velocimetry, and in computer vision applications, such as image compression and 3D shape reconstruction [15]. Cross-correlation relies on strong image textures; thus, DIC is often enhanced with painting or powder coating of macroscopic samples [7]. While this is rarely feasible for biological samples, DIC has still been an accurate method for full-field displacement estimation, especially in samples that have intrinsically high textural contrast [6, 7]. However, because of the requirement to support sub-pixel accuracy and local deformation, the iterative and subset-based search methods employed in DIC can take minutes for image pairs, and hours or days for videos.

A fast approximation of displacement between two images can be achieved using optical flow (OF), which typically estimates velocity from local spatiotemporal gradients of the image intensity, which must be combined with regularisation, such as of local smoothness, to converge to a unique solution [16]. The development of such methods is an active field of research, with multiple benchmarking efforts in place to keep track of improvements [17, 18]. For example, the Middlebury optical flow challenge currently lists 190 competing methods [17]. Traditional OF methods predominantly rely on the assumption of ‘brightness constancy’, i.e., that the intensity of the imaged scene does not change, but merely shifts locally in the field of view. OF further struggles with large displacements that exceed the spatial resolution of the moving objects. Thus, OF methods typically excel in rigid image motion with strong image features and gradients, such as in self-driving cars or video compression [18]. However, OF performs poorly in biological images, which typically have lower textural contrast, signal-to-noise ratio and fluctuating intensities. Because of this, DIC is often preferred for biomechanics analyses [6, 7]. Thus, there is a present need for accessible methods that can combine both high accuracy and computational speed.

Deep learning has the potential to address this challenge with a feed-forward scheme that could exceed the speeds of even optical flow methods. A major challenge in this task is that, in most scenarios, ground truths measurements of displacement are unavailable. Because of this, DIC and OF methods are typically evaluated based on some benchmark dataset [19] with simulated or model-based approximations of displacements, and are assumed to readily extend to other types of image data. Thus, appropriate metrics and training data consistent with biological images must be generated for any deep learning task. Deep learning has been applied to OF for computer vision, particularly, in simulated and animated graphics [20]. Methods such as the milestone FlowNet [21] achieve powerful OF performance by training on artificial scenes, typically via publicly available datasets. Additionally, several recent works have investigated the use of deep learning for DIC motivated by material testing. Boukhtache et al. [22] have presented StrainNet, a model based on tracking subpixel displacement between images of speckle-painted materials, for instance, in tensile testing. While the StrainNet network was based on the FlowNet architecture, it was instead trained on synthetic images of speckle and a grid-based displacement simulation. The model performed well in estimating microscale deformation of solid material. Yang et al. [23] have extended StrainNet by end-to-end training directly on local strain, enhancing strain sensitivity. Full-field displacement estimation using deep learning, however, has yet to be developed for biological images, which distinctly have a broad range of image features, contrasts and sources of noise. As shown in this work, these properties makes StrainNet and similar materials-testing-inspired networks poorly suited to the task.

In this Paper, we demonstrate an experimentally unsupervised deep learning method for estimating full-field displacement in biological images: meaning that this method can be applied to any biological image pair or video with no prior knowledge of displacement or additional experiments. We refer to our network as: Displacement Estimation FOR Microscopy–’DEFORM-Net’. DEFORM-Net combines images from different modalities, and generates synthetic supervised data consistent with typical power spectral characteristics of biological deformation using fractal Perlin noise, as well as system noise processes that, by design, invalidate the brightness constancy assumptions. We explore various similarity metrics, including perception-based losses, as well as an image correlation based metric using a forward simulation of deformation to improve performance and network generality. DEFORM-Net is evaluated on real microscopy videos of beating mouse cardiomyocytes [24, 25] and pulsating larval epithelial cells (LECs) in *Drosophila* pupae [26]. We show that DEFORM-Net can match and exceed the accuracy of DIC and the speed of OF, with near video-rate inference speed and high spatial sampling. To assist rapid iteration and testing, we provide DEFORM-Net as open source, including a standalone compiled version for inference and integration with the ImageJ/FIJI platform [27]. Our method is readily adaptable and will empower powerful quantification for dynamic bioimaging challenges.

## 2 Results

### 2.1 Full-field displacement estimation with deep learning

The problem of estimating full-field displacements from image pairs is illustrated in Fig. 1(a–d). This example shows false-coloured widefield fluorescence microscopy images of two consecutive frames, termed as reference (Ref.) and deformed (Def.), from a video of a cluster of spontaneously beating neonatal mouse cardiomyocyte cells. The displacement field between the reference and deformed states has horizontal and vertical (or *x* and *y*) components that can be visualised in a vector form, **u** = (*u*_*x*_, *u*_*y*_), (Fig. 1(c)) or in a phasor form, where *A* = |**u**| and *ϕ* = ∠ **u**, (Fig. 1(d)). The phasor visualisation encodes the displacement magnitude as the value (V) and the angle as the hue (H) in the hue-saturation-value (HSV) colour space. Figure 1(e) shows how this phasor representation can be a convenient means of illustrating the displacement field with a high spatial resolution.

**Figure 1.**
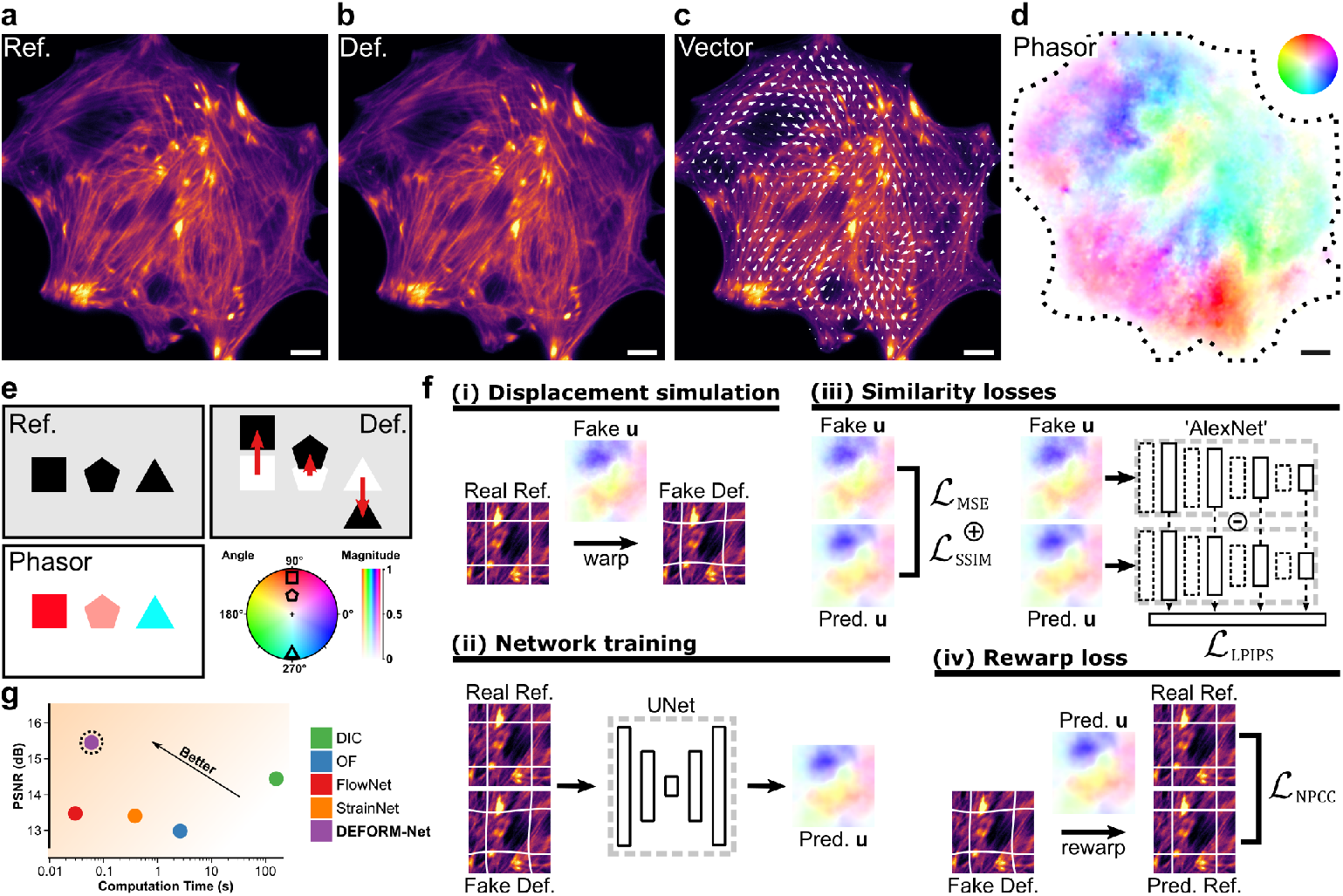
Illustration of full-field displacement estimation. (a) Reference (Ref.) and (b) deformed (Def.) fluorescence microscopy images of mouse cardiomyocytes. The full-field displacement from the reference to the deformed state can be represented by a (c) vector image or (d) phasor plot, where the hue represents the angle and saturation represents the magnitude of the displacement vector. Scale bars are 10 *μ*m. (e) Visual demonstration of the mapping from displacement to colour. (f) Graphical representation of the (i, ii) deep learning method and (iii, iv) loss functions, where **u** is the displacement vector field and the terms Real and Fake refer, respectively, to real microscopy data and simulated data. (g) Comparison of the performance displacement estimation methods in real data quantified using the peak signal-to-noise ratio (PSNR).

DEFORM-Net comprises several key components that enable it to estimate an accurate displacement field between input image pairs (Fig. 1(f)). Because ground truths are unavailable in this problem, we simulate realistic displacement fields (Fake **u**) and use them to generate synthetic pairs of reference (Real Ref.) and deformed (Fake Def.) images for supervised training (Fig. 1(f)(i)). These images are further augmented with noise processes that mimic those in microscopy experiments. This is detailed in the Methods section and evaluated in the later Results sections. We train an artificial convolutional neural network, based on a U-Net architecture, to transform the image pairs into a predicted displacement field (Pred. **u**) using several training loss functions (Fig. 1(f)(ii)). First, we implement conventional pixel-wise similarity losses between the predicted displacement field and the simulated ground truth, including the mean-squared error (MSE) and the structured similarity index metric (SSIM) (Fig. 1(f)(iii)). SSIM is a common metric that emphasises matching of perceived textures between image patches and helps counteract the blurring effects associated with absolute error-based estimates [28]. We enhance this with a learned perceptual similarity loss (LPIPS), which is emerging as a powerful image similarity metric based on intercepting intermediate feature maps from pre-trained large-scale classification networks (here, the AlexNet) [29] (Fig. 1(f)(iii)). We evaluated larger-scale networks for LPIPS, such as the VGG-19 model, however, they have not resulted in significantly different or improved performance, which was similar to the findings of the original work [29]. Finally, we include an unsupervised loss metric based on image correlation that we term as ‘rewarp loss’ (Fig. 1(f)(iv)). The synthetic deformed image (Fake Def.) is ‘undeformed’ using the predicted displacement field (Pred. **u**), generating an estimate of the real reference image (Pred. Ref.). This is then compared to the real reference image using negative Pearson correlation coefficient (NPCC). As such, no knowledge of the ground truth is required to calculate the rewarp loss. This methodology leads to improved performance in real biological videos compared to previous methods (Fig. 1(g)). We describe these performance values and justify the methodology in the following Results sections, first with simulated supervised test data and then with real experimental data.

### 2.2 Simulating biologically realistic displacements

A major challenge in network training is the availability of accurate ground truth data. While OF networks, such as FlowNet, employ animated scenes and DIC networks, such as StrainNet, employ the simulation of speckle, we demonstrate that these are insufficient at capturing displacements and image content of biological images, leading to poor performance. Figures 2(a) and (b) show representative reference frames from widefield fluorescence microscopy of mouse cardiomyocytes (SiR-actin) and confocal microscopy of *Drosophila* LECs (F-actin, GMA-GFP), respectively, with the associated displacement fields estimated using conventional DIC and OF methods between the reference frame and the next recorded frame. The implementation details of DIC and OF, along with information about the biological samples, are in the Methods Section 5.7. We can see that the estimated displacement comprises large scale shifts in various areas of the images, as well as a range of higher spatial frequency variations, both in amplitude and direction. The DIC estimates are of lower spatial resolution to that of OF due to the finite subset size, but are typically regarded as more accurate [15]. While both displacement fields are scaled for visualisation, the maximum displacement magnitude was approximately 200 nm (2 pixels) in the cardiomyocyte and 1.7 *μ*m (8 pixels) in the *Drosophila* data. The larger displacements in the *Drosophila* data led to a discrepancy between DIC and OF estimates, likely due to the well-established problem of OF with large displacements [16].

**Figure 2.**
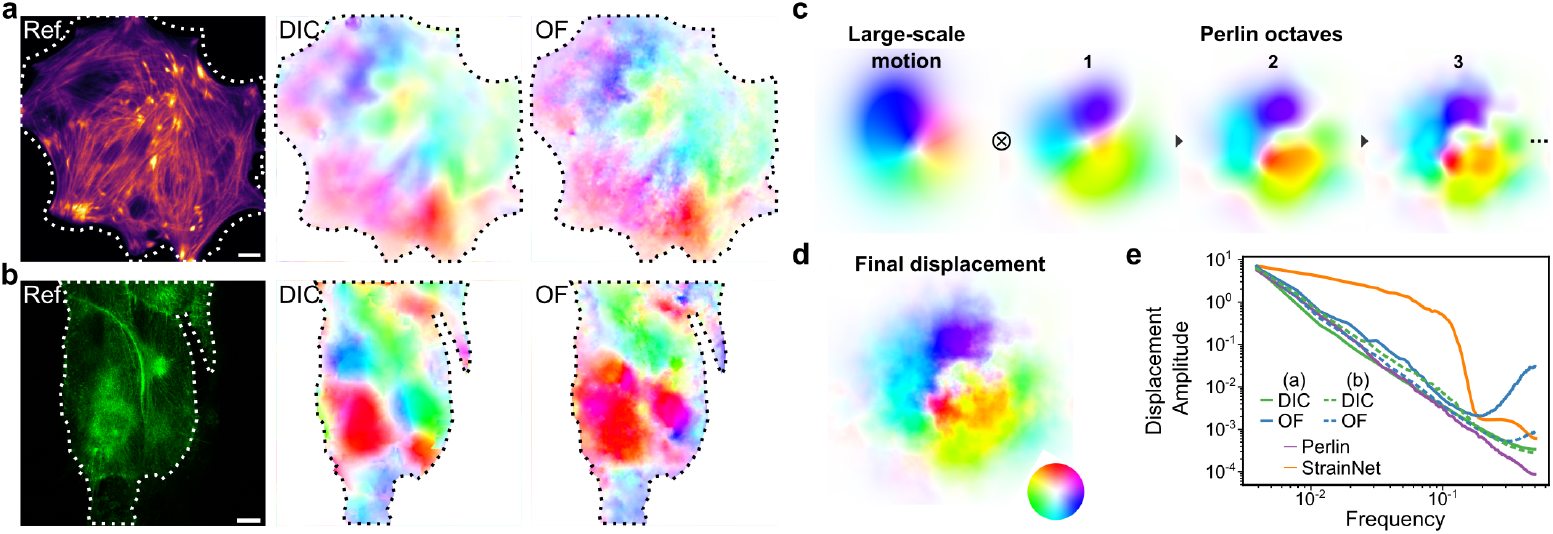
Simulation of realistic displacement fields. (a, b) Reference microscopy images of (a) mouse cardiomyocytes and (b) *Drosophila* LECs with representative displacement fields estimated using DIC and OF methods. Scale bars are 10 *μ*m. (c) Displacement vector field simulation by multiplying random Gaussian large-scale motion with progressively refined fractal Perlin noise. (d) Final simulated random displacement field with controllable spatial power-spectral density. (e) The spatial power-spectral density of displacement amplitudes estimated using DIC and OF from data in (a) and (b) compared to simulated data using Perlin and StrainNet [22] methods.

To capture such properties of biological motion, we propose displacement simulation based on a random fractal Perlin noise process [30]. This is detailed in the Methods Section 5.1; however, briefly, we first generate large-scale motion using several randomly positioned Gaussian peaks with a random displacement phasor (Fig. 2c)). We then multiply this with several ‘octaves’ of random Perlin noise (amplitude and direction), with each octave possessing progressively refined spatial resolution and reduced energy to generate the final displacement field (Fig. 2(d)). Perlin noise is a procedural noise originally developed as a method to generate realistic image textures [30]. What makes this noise well suited to the task is the precise control of the spatial power spectral density (PSD) of the generated fields, i.e., the energy associated with a particular spatial frequency. Figure 2(e) shows the mean PSD across (n=50) DIC and OF estimates in biological samples in Figs. 2(a) and (b). We immediately see a linear log-log decay of displacement amplitude with spatial frequency consistent between samples and DIC and OF methods. This property of the PSD is often found in natural images, and exploited as an image prior in many reconstruction methods [31]. Our Perlin-based method naturally satisfies this PSD shape. This is an advantage to the grid-based generation method employed in StrainNet [22], which inevitably leads to a hard cut-off response. The inflection in PSD seen in the OF estimates is likely due to the instability in the OF algorithm, which arises when the total displacement exceeds its spatial frequency [16]. We do not compare to learned OF methods, as their training is based on computer-generated scenes [17, 18].

### 2.3 DEFORM-Net outperforms other methods in simulated data

We train DEFORM-Net using simulated training data, generated using reference image frames from four datasets, each acquired with a different microscopy imaging modality (Figs. 3(a–d)). These are: widefield fluorescence microscopy, brightfield microscopy and differential interference contrast microscopy (DICM) of mouse cardiomyocytes, and confocal fluorescence microscopy of LECs of a *Drosophila* pupa with labelled F-actin (GMA-GFP). Specifically, we generate 2,337 frame pairs, including a real reference image, a simulated deformed image and the associated simulated displacement field, from randomly cropped regions of interest from each frame of the datasets. These regions are augmented with random rotation and image transpose operations. After the complete training set was generated, it was randomly split into 1,825 training and 256 validation pairs, as well as 256 frame pairs exclusively used for testing. The training pairs were used for evaluating the loss for back-propagation, while the validation pairs were used to monitor for convergence and overfitting. The testing pairs were not used in the network training process, and were only used to quantify performance. All image modalities were used to train a combined model for evaluation in the first instance. While evaluating performance in simulated images is not representative of ultimate performance in real data, we use this for an initial comparison of existing methods.

**Figure 3.**
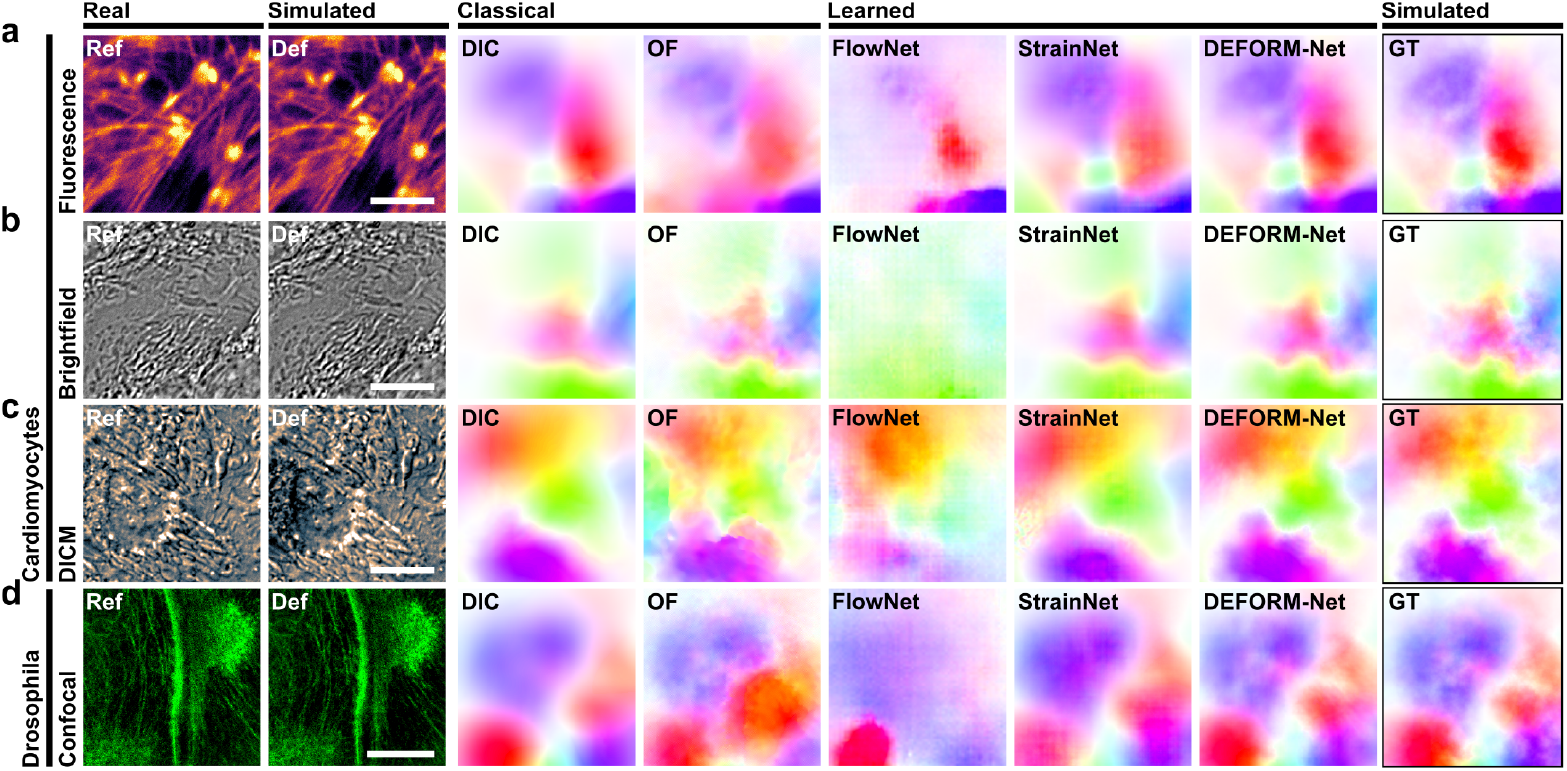
Visual comparison of methods in different imaging modalities and contrasts in simulated data. (a–d) Representative microscopy images and estimated displacement fields of (a–c) mouse cardiomyocytes using widefield fluorescence microscopy, brightfield microscopy and differential interference contrast microscopy (DICM), and (d) *Drosophila* LECs using confocal fluorescence microscopy (F-actin, GMA-GFP). Scale bars are 10 *μ*m.

Figure 3 compares the performance of classical and learned methods in a representative image randomly selected from the testing set for each modality. Specifically, we perform conventional DIC using ‘ncorr2’ software [32] and OF using a pyramidal TV-L1 solver [33], which were chosen because they output the highest accuracy estimates compared to other common open-source methods for this data (see Methods Section 5.5). We compare this to learned FlowNet and StrainNet methods. The original StrainNet network (trained on artificial speckle) performed poorly; thus, for a fair comparison, we have retrained StrainNet with the images used in this work, but simulated with the grid-based approach scaled to the higher displacement amplitudes seen in the images. Because StrainNet relies on separated generation of displacements at various resolutions, the training set expanded to over 20,000 image pairs and was trained for 6 ⨯longer. FlowNet, however, was not retrained as it relies on the use of animated graphics datasets. Like conventional OF, it is expected that it can be applied indiscriminately to all data.

Visually comparing the different methods to the ground truth (GT), we see that DIC outputs an accurate but substantially blurred estimate. OF, however, produces higher spatial resolution outputs with several sections in the images that fail to match the GT. FlowNet performs overall poorly, especially in images with low image contrast and high noise (Figs. 3(b) and (d)). While StrainNet estimates are adequate compared to DIC, DEFORM-Net substantially improves the estimate quality, especially in terms of spatial resolution. Notably, StrainNet has directly implemented the FlowNet network architecture (FlowNetF), which included a multiscale loss in the decoder side. We found that this architecture resulted in similar loss values to a conventional U-Net; however, it required four times longer to train.

We assess performance through various error metrics (Tab. 1) based on the training loss functions described in Section 2.1. Specifically, we use peak signal-to-noise ratio (PSNR) as 10 · log_10_(1*/*MSE), such that a higher value corresponds to better performance for all metrics. Notably, as it is a log-scale metric, a 3-dB variation in PSNR corresponds to a two-fold variation in accuracy. Further, we scale the metrics to the range of 0 to 100 for convenient comparison. DEFORM-Net outperforms all other methods on all loss metrics, including DIC. Importantly, the inference time per frame is substantially improved over classical methods. Much faster OF methods could be implemented; however, the slower pyramidal and iterative method was required to achieve satisfactory performance in biological images. While FlowNet is the fastest, it evaluates a 4*×* sub-sampled output compared to other methods; StrainNet utilises an identical inference graph to FlowNet without subsampling, for a more direct comparison. All learned methods, however, breach the limiting two-minute processing time of DIC.

**Table 1.**
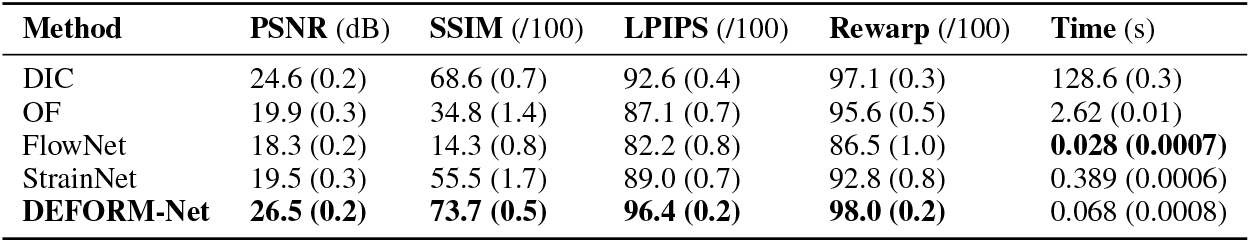
Performance statistics of displacement estimation methods in simulated data. Data is presented as mean value and standard error of mean in brackets (*n* = 50). Highest performance values within standard error bounds are highlighted in bold. PSNR: peak signal-to-noise ratio of MSE; SSIM: structured similarity index metric; LPIPS: learned perceptual image patch similarity; Rewarp: Pearson correlation coefficient of rewarp loss.

### 2.4 DEFORM-Net outperforms other methods in real microscopy data

We next evaluate DEFORM-Net on real experimental data. Because there is no access to independent ground truth data, we consider an independent method of validation by manually tracking the displacement of local features using the ImageJ manual tracking functionality. Whilst the rewarp loss could be used as an unsupervised metric of performance, we found that it suffers from local minima, thus, making it unsuitable for accurate quantification of error (Supplementary Note S1). First, we have manually generated 9 sets of trajectories that are visually accurate from the raw confocal microscopy videos of *Drosophila* LECs with fluorescently labelled F-actin (GMA-GFP) (Fig. 4(a)). We note that the raw image frames were used in training DEFORM-Net using simulated data, however, the real motion was never seen by the training. Each trajectory comprises 87 coordinate points corresponding to each consecutive frame pair in the entire video. We then compare the displacement with that of the corresponding estimate in the nearest pixel for each method. Figure 4(c) shows a representative displacement track generated by conventional methods compared to DEFORM-Net. We then evaluate the MSE of their distance metric, and the MSE for each frame and track are averaged and converted to the log-scale PSNR for comparison (Fig. 4(e)). We see that the performance in real data is substantially higher for DIC and DEFORM-Net compared to other methods, as expected from the simulated tests. We use these tracks and the manual-tracking-based metric for further evaluations.

**Figure 4.**
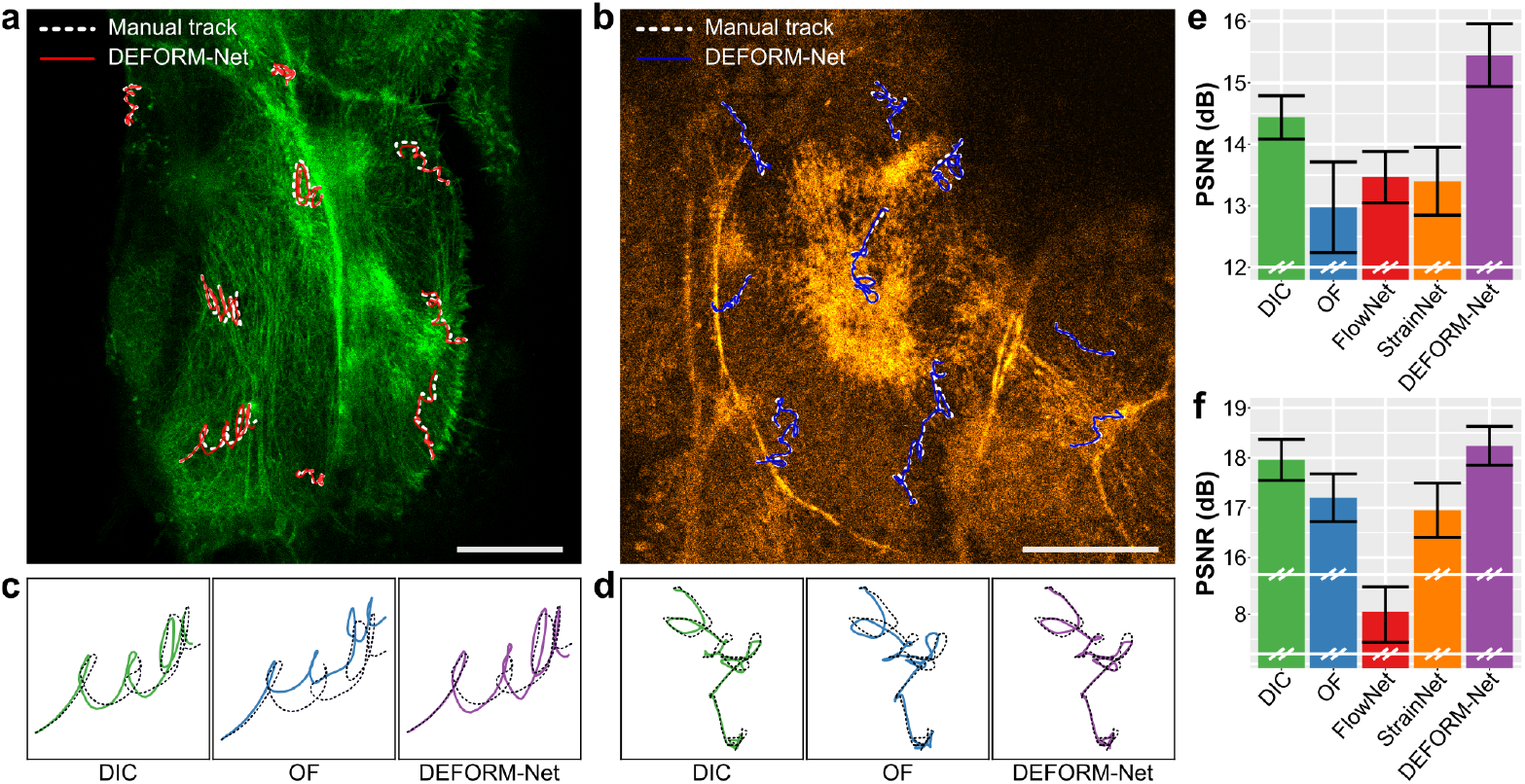
Performance evaluation in real displacement data. Visualisation of manually tracked displacements in *Drosophila* confocal microscopy videos of (a) F-actin, GMA-GFP (used in the training set) and (b) myosin, Sqh::GFP (unseen by training). Scale bars are 20 *μ*m. Representative displacement tracks in (c) F-actin and (d) myosin datasets evaluated using conventional methods compared to DEFORM-Net. Displacement error compared to manual tracking in (e) F-actin (n = 9) and (f) myosin (n = 9) datasets. Error bars are standard error of mean.

To demonstrate the generalisation of DEFORM-Net, i.e., its ability to perform in new, unseen data, we evaluate its performance on a different confocal microscopy video of *Drosophila* LECs, however, now expressing fluorescently labelled Myosin II regulatory light chain (Sgh::GFP) [26]. While Sqh::GFP, similar to GMA-GFP, is visualising the pulsed contractions of the LECs, both this sample and the source of contrast have not been seen by the network training process (cross-domain inference). As for GMA-GFP, 9 sets of manual trajectories were generated for Sqh::GFP in ImageJ (Fig. 4(b)). Figure 4(d) shows a representative trajectory using conventional methods, while Figure 4(f) provides detailed mean PSNR of the distance error metric. We can see that again both DEFORM-Net and DIC outperform other methods. Interestingly, FlowNet was unable to estimate meaningful displacements in this data. This is likely due to the substantially higher background noise in the data. Because of this, the parameters of DIC used for all previous data were insufficient in estimating accurate displacements. Thus, we have expanded the spacing and subset ranges from 3 and 30, to 5 and 50 pixels, respectively. For all methods, we have further smoothed the input images by a Gaussian blur with a sigma of 1 pixel. These results indicate that even without retraining, DEFORM-Net can generalise to other microscopy data. We investigate this further in the next section.

### 2.5 Including microscopy-informed noise in training data improves generalisation

A key part of the performance of DEFORM-Net is the added treatment of image intensity noise. Compared to natural macroscopic scenes, biological images are often challenged by many sources of noise, such as from poor biological contrast, background noise or poor sensitivity of instruments. Beyond random noise that follows a combination of a Poisson and Gaussian process, which are added to our training data, additional large-scale fluctuations in intensity can invalidate the brightness constancy assumption in OF methods.

Figures 5(a) and (b) illustrate this issue in images of *Drosophila* LECs. The DIC vector image shows a pattern of pulsatile motion in two LECs. The red circle marks an area of converging displacements, and the red band highlights an area where an optical occlusion (a fold in the pupal cuticle) dims the fluorescence signal captured by the confocal scan. The OF estimate, however, shows a different direction in the displacement vectors. Compared to manual tracking, which we use as the baseline, DIC estimates are more accurate (Fig. 5(c)). The poor performance of OF is likely because the intensity between frames is not consistent. In fact, this is common to fluorescence imaging due to photobleaching, fluctuation in the fluorescent background, and variations in the absorption and scattering of light in a living tissue.

**Figure 5.**
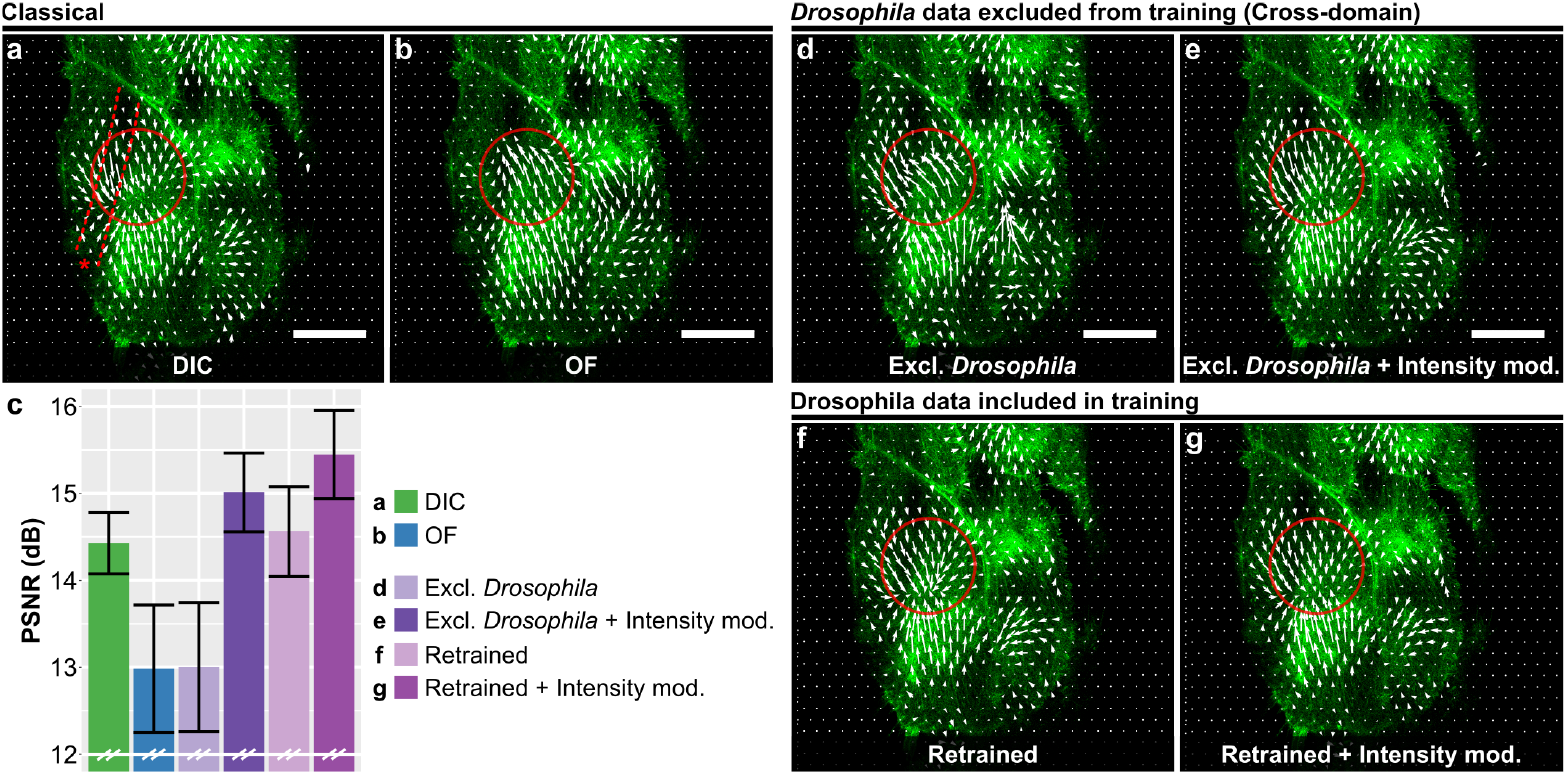
The role of noise and retraining in generalisation of network performance. (a–b) Displacement vectors from a single frame pair from a video of LECs in a *Drosophila* pupa evaluated using conventional (a) DIC and (b) OF methods. (c) Accuracy of methods compared to manual tracking. Error bars are standard error of mean. (d–g) Displacement vectors from networks trained using different methods: (d) cardiomyocyte data only (excluding *Drosophila* data); (e) cardiomyocyte data only, with intensity modulation noise; (f) retrained on *Drosophila* data; and, (g) retrained on *Drosophila* data with intensity modulation noise. Scale bars are 20 *μ*m.

Naively, we would like DEFORM-Net to be applicable to all images with a use-case similar to that of DIC or OF, i.e., our network should demonstrate cross-domain performance. As such, we train a DEFORM-Net model with data comprising solely the multimodal cardiomyocyte images, specifically excluding any *Drosophila* images, and with only added Poisson and Gaussian noise. However, Fig. 5(d) shows that the output of such a network exhibits similar artefacts as that in OF, suggesting that this network learns some similar properties of brightness constancy to OF models. To overcome this, we intentionally violate brightness constancy by introducing another noise, which we term as ‘intensity modulation’ noise. Specifically, we introduce additive large-scale and low-spatial-resolution noise, also modelled using our Perlin noise process (but with fewer octaves), with a maximum peak size of 10% of the image dynamic range. When trained with such data, even excluding any *Drosophila* images, the performance is substantially enhanced and exceeds that of DIC (Fig. 5(e)).

By retraining DEFORM-Net with data that includes the *Drosophila* images, the overall performance increases substantially for the Poisson and Gaussian noise-only case (Fig. 5(f)). However, retraining only has a marginal improvement when intensity modulation noise is included (Fig. 5(g)). This performance (Fig. 5(c)) suggests that intensity modulation noise is important for enhancing network generalisation, perhaps more so than the breadth of training images, and has thus been implemented in all other evaluations in this work. Our combined loss functions further contribute to this generalisable performance. We detail these losses and perform an ablation study in Supplementary Note S2. We further evaluate the performance and generalisation on publicly available data provided by Lye et al. [34] in Supplementary Note S3. There we see similar improvements to performance with the addition of ‘intensity modulation’ noise during training. While ‘intensity modulation’ may not be critical to all biological samples and modalities, the results suggest that it is a useful tunable parameter for training a robust network.

### 2.6 Demonstration of DEFORM-Net in biological videos

Figure 6(a) shows spontaneous synchronised beating of cardiomyocyte cells, which was evaluated as the mean of the displacement amplitude in the field of view over the 15-second video. Examples of displacement fields associated with the primary and secondary peaks of the beat are visualised in the insets in Fig. 6(a)(i) and (ii), respectively, revealing a pattern of simultaneous contraction succeeded by smaller relaxations. Studying the spontaneous beating of neonatal cardiomyocytes and the intercellular communication between neighbouring cells provides invaluable insights into cardiac activity, intrinsic pacemaker activity, and developmental signalling pathways [35]. Understanding these fundamental mechanisms not only elucidates the normal functioning of the developing heart but also offers critical avenues for research into various diseases and potential therapeutic interventions. This data was recorded with DCIM, and individual image frames were used as part of the simulated training data. This represents a use case where the acquired microscopy data is directly used to train and optimise a bespoke DEFORM-Net model. The full video is available as a Supplementary Movie S1.

**Figure 6.**
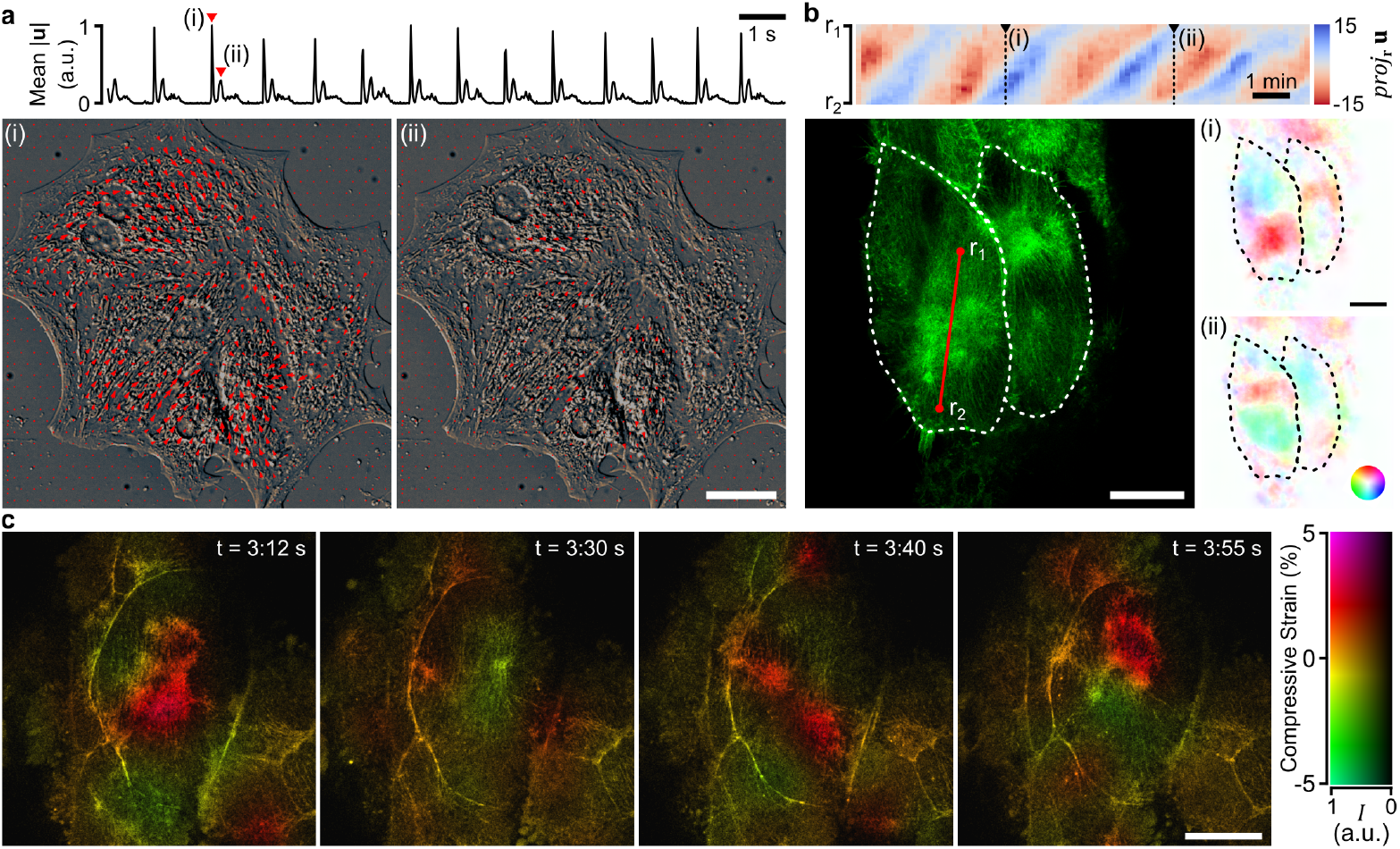
Demonstration of full-field displacement estimation. (a) Beating mouse cardiomyocytes imaged with DICM show a regular beating pace from the mean displacement magnitude across frames. Snapshots of displacement field vectors are from (a)(i) primary and (a)(ii) secondary peaks. (b) Displacement projected along the *r*_1_*r*_2_ vector from a video of two pulsating *Drosophila* LECs (confocal microscopy, F-actin, GMA-GFP) marked by the dashed white line. Insets show the displacement phasor maps at frames (b)(i) and (b)(ii). (c) Local compressive strain maps (−*dV/V*) false-coloured as hue shifts in image frames of a constricting *Drosophila* LEC (confocal microscopy, myosin, Sqh::GFP), unseen by network training. Scale bars are 20 *μ*m.

Figure 6(b) shows the pulsatile motion of the F-actin cytoskeleton (GMA-GFP) in *Drosophila* LECs as a projection of the displacement field along the line indicated by points *r*_1_ and *r*_2_. This line plots a trajectory of wave-like motion along the cell’s long axis. Insets in Fig. 6(b)(i) and (ii) show displacement phasor plots from two time points. Such pulsed contractions are important in *Drosophila* development because they are involved in the apical constriction of LECs, which precedes their programmed cell death during metamorphosis, when larval tissues are replaced by adult tissues [26]. Understanding pulsed contractions will provide mechanistic insights into congenital conditions caused by defective contractility, for example, neural tube disorders, such as spina bifida [36]. The full phasor and vector videos are available as a Supplementary Movies S2 and S3.

For the demonstration in Fig. 6(b), we have used the DEFORM-Net model that was trained excluding all images of *Drosophila* LECs from Section 2.5. While retraining the model provides some marginal improvement, these results demonstrate that our network can generalise to unseen data from a different imaging modality and with different image contrast. Figure 6(c) further supports this by demonstrating displacement-derived local compressive strain maps in frames from a video of a constricting *Drosophila* LEC with labelled myosin (Sgh::GFP). Compressive strain represents the relative change in volume, which was evaluated as the local divergence of the displacement field, i.e., *dV/V*_0_ = *∂u*_*x*_*/∂x* + *∂u*_*y*_*/∂y*. This value was inverted to relate a positive value with compression. The fluorescence labels the Myosin II regulatory light chain, which exhibits coalescence (increase in local protein density) associated with local contraction of the cytoskeleton. In fact, local intensities were previously used as a metric of contraction [26]. In Fig. 6(c), we see a correspondence in local compressive strain with the coalescence of myosin, supporting previous observations but not with direct quantitative data. The full strain and vector videos are available as a Supplementary Movies S4 and S5. Importantly, this video was never used in training any DEFORM-Net model, nor was it used to optimise the displacement simulations or loss functions.

## 3 Discussion

DEFORM-Net is a powerful method for accurate and rapid estimation of displacement in biological images that combines the accuracy of DIC with rapid and high-resolution feed-forward inference of deep learning. Conventional DIC and OF methods are routinely used for advanced biological analyses [7, 6, 8, 9, 10, 11, 12]. However, the accuracy of such estimates is rarely validated independently. This challenge is compounded by the fact the accuracy is sample and contrast dependent. Because of this, the general wisdom suggests that one should select a method that produces outputs that are consistent with the motion intuitively observed in the images. In the same way, we foresee that DEFORM-Net could be used with some selection of pre-processing and the model. We have evidenced that DEFORM-Net can perform well in generalisation with limited data, both with data that was never seen by the training and user optimisation process (Fig. 4), and by specifically excluding data from model training (Fig. 5). We note that in the latter case, the model has not seen any *Drosophila* images nor any confocal microscopy scans. If retraining is required, such experimentally unsupervised datasets are reasonable to acquire in a single or few sample scans, e.g., solely the data that is desired to be analysed. This suggests that no additional experimental time may be required to train the model. Further, one may combine existing data with new sample frames for combined multi-domain training, as we have done in this work.

To facilitate rapid dissemination, we provide DEFORM-Net as open source (see Methods section). Because training new models can be a challenge both in the requirement for advanced hardware and expertise, we also provide the capacity to perform rapid inference with pre-trained models through a compiled executable through a command-line interface or using the ImageJ/FIJI platform [27]. For this, we further make available several key models from this paper and the associated datasets. Additional details on the software is included in the Supplementary Information. We hope that such open-source deposition of our method facilitates the evaluation of DEFORM-Net with various microscopy modalities, biological systems and contrasts. DEFORM-Net should be evaluated for each type of biological contrast and dynamics, with appropriate tuning of the loss functions and training parameters, for optimal performance. For instance this, would be particularly beneficial to high-content and high-throughput systems, which can avail of the rapid inference speed of deep learning.

DEFORM-Net may also be applied to multimodal image data. For instance, naively, DEFORM-Net can evaluate motion independently between each individual image channel. The estimated displacement fields may then be averaged together. Alternatively, DEFORM-Net can be retrained to accept a composite image with a specified number of channels; however, DEFORM-Net would have to be retrained for each particular data and channel orientation.

The fractal Perlin noise simulation of displacement fields is a critical part of DEFORM-Net and its improved performance compared to StrainNet and FlowNet methods. The PSD of the simulated displacement fields can be tailored to that expected in the real data. However, further improvements may be gained by introducing some PSD regularisation in the training loss functions, which can be important priors in biological data [37]. Specifically, the PSD of the network output can be constrained to match that of natural images [31]. The rewarp and LPIPS losses further provide options for more natural and physics-based regularisation of network training. Rewarp loss is not globally optimisable because of local minima, especially at higher spatial resolutions (Supplementary Note S1). In DIC, such metrics are regularised by analysing patches and constraining the local deformation to a set of affine transforms [32]. While this could be adapted to the loss function, its evaluation would be substantially slowed as this is the critically slow component of the DIC method.

The field of learned OF methods is flourishing with many new advances that can be investigated for DEFORM-Net. While we have not exhaustively compared our network performance across this field, FlowNet still remains as an important recent benchmark with only marginally poorer performance to the current leading methods. However, in our work we have observed that such methods are ill-suited to the domain of biological images because of the routine violation of the brightness constancy assumption and the typically high background noise. For instance, RAFT [38], a previous leader in learned OF, did not outperform FlowNet in these images. Despite this, the major advances in learned optical flow could be used as inspiration to enhance DEFORM-Net. Specifically, well-performing network architectures could be evaluated using the DEFORM-Net data generation and training scheme in future advances.

Perceptual networks and the derived loss functions are an exceptionally interesting prospect for displacement estimation [29]. Qualitatively, human observation can track a greater range of motion between overlapped image frames, even in areas where DIC and OF fail, likely utilising a breadth of image contrast, texture and feature identification abilities. Thus, displacement estimation tasks could be enhanced by transforming input data to new perception-based representations, or by combining several networks that excel at complementary tasks. However, human perception is challenged by many artefacts where motion is inferred where no motion might exist [39], which may lead to fascinating mergers of cognition-based networks with physics-based systems.

## 4 Conclusion

We have presented DEFORM-Net, a deep-learning method for estimating full-field displacement between biological images or video frames. DEFORM-Net outperforms conventional digital image correlation and optical flow methods, as well as their learned StrainNet and FlowNet counterparts, in microscopy data that is typically challenged by poor sample signal, low contrast and background noise. This performance was afforded by the random fractal Perlin-noise simulation with a controlled power-spectral density matching natural displacements, as well as the addition of intensity noise and several training loss functions. The trained models exhibited good performance on videos of beating mouse cardiomyocytes and pulsating *Drosophila* cells, including in cross-domain scenarios, suggesting good generalisation. We present DEFORM-Net as open source with inference implemented in the ImageJ/FIJI platform, towards rapid evaluation across many applied problems in biology and medicine.

## 5 Methods

### 5.1 Simulating full-field displacement with fractal Perlin noise

The simulated displacement fields are generated in two stages. First, several two-dimensional Gaussian peaks are placed at randomised locations and combined additively. Specifically, we generate six peaks, while their location, magnitude and standard deviation are randomly sampled from independent Gaussian distributions.

Second, the large-scale displacements are combined with fractal Perlin noise [30], a type of smooth pseudo-random pattern, the appearance of which is often likened to clouds. This introduces small-scale variations which closely match the displacement distributions observed within our microscopy images. Conveniently, we utilise the open `perlin_numpy` [40] package for this. Briefly, the generation procedure for fractal Perlin noise is as follows: a grid is defined, the scale of which determines the scale of the variations in the noise. A random unit vector, known as the gradient vector, is generated at each grid node. For each point in the field, the four closest grid nodes are considered. The dot product between each node’s gradient vector and the displacement vector of the point from the node is calculated. Then, the final noise value is obtained by smoothly interpolating between these four results, based upon the distance between the point and each node. The interpolation function used should have stationary points at the grid nodes, as this leads to the noise values equalling zero at each node, binding the scale of noise variations to the grid scale. Fractal noise is created by adding many layers of Perlin noise, referred to as ‘octaves’. Each octave’s magnitude and grid spacing are scaled down from the previous by factors known as the ‘persistence’ and ‘lacunarity’, respectively.

The parameters used for large-scale noise and fractal Perlin noise generation were chosen based on a visual match to the DIC and OF estimates in biological data. The exact values may be inspected in the `perlin_generator` module of the supporting open-source code (See Methods Section 5.4).

### 5.2 Generation of supervised training data

Training DEFORM-Net requires input images in the reference and deformed configuration, as well as the corresponding displacement field to evaluate the supervised training losses. To generate this set of data, we used individual frames from real microscopy videos. For each image pair, a randomly cropped area (here, 512*×*512 pix) from a real frame was used as the reference image, with the corresponding grid spatial coordinates, *I*_*R*_(*x, y*). Then a simulated displacement field, 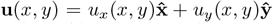, is used to deform the reference image to a simulated deformed image, *I*_*D*_. This is achieved by evaluating the reference image at new transformed coordinates: (*x*^*′*^, *y*^*′*^) = (*x*−*u*_*x*_, *y*−*u*_*y*_), i.e., *I*_*D*_(*x, y*) = *I*_*R*_(*x*^*′*^, *y*^*′*^). Because (*x*^*′*^, *y*^*′*^) are subpixel coordinates, we used two-dimensional interpolation based on a bivariate spline approximation over a rectangular mesh. This procedure was repeated for each image frame in the real microscopy videos, and the final data was randomly shuffled and partitioned into exclusive training, validation and testing subsets (exact image numbers are provided in the relevant Results sections). The training data was used for training, loss evaluation and backpropagation, while the validation data was used to visualise convergence. Testing data was used for all evaluations in the results with no influence on the training process.

### 5.3 Network architecture and loss functions

DEFORM-Net uses a 7-level U-Net autoencoder [41], which outputs full-resolution *W×H×*2 displacement estimates (one for the *x* and *y* displacement component), where *W* is the number of pixels in *x* direction and *H* is the number of pixels in *y*−direction. The reference and deformed frames are concatenated and input as a *W×H ×*6 tensor. This is to support 3-channel RGB colour images; however, greyscale images were used in this instance because all biological data were single-intensity images. The number of input channels does not appreciably increase inference or training times, as the first convolutional layer transforms the input to a set 64 channels regardless.

DEFORM-Net was trained using a desktop computer equipped with an NVIDIA GTX 1080Ti GPU and an Intel Xeon W-2145 CPU. Training took approximately 5 hours for 300 epochs and 1,825 frame pairs with 512 *×* 512 pixels each.

A combination of supervised training loss functions were used to train our final model, each based upon comparing the estimated displacement fields to the simulated ground truth, including: (1) the Mean Squared Error (MSE), (2) the Structural Similarity Index Measure (SSIM) [28], and (3) the Learned Perceptual Image Patch Similarity (LPIPS) [29]. In addition to these, (4) an unsupervised ‘rewarp’ loss was used.

(1) MSE loss evaluates the mean squared error of the Euclidean distance metric between the network output and the ground truth:

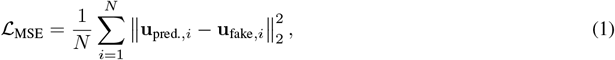

where **u**_pred._ and **u**_fake_ are the network outputs (predicted) and simulated (fake) displacement fields, respectively (Fig. 1(f)). This loss is typical in many deep learning-based displacement estimation methods [21, 22, 42], despite not accurately reflecting several critical aspects of image quality, such as the level of blurring or noise [43]. The MSE is also commonly referred to as the endpoint error (EPE) when evaluated on vector values, such as displacements.

(2) SSIM is frequently used as a metric designed to perceptually compare larger scale features within images [28, 43]. It aims to capture structural differences, such as those arising from blurring or noise, independent of changes in image brightness (which represents displacement magnitude in our case). This is achieved by splitting the input images into patches for comparison (here, 11-pixels wide), rather than only considering corresponding individual pixels. The SSIM is most commonly written as a product of three components:

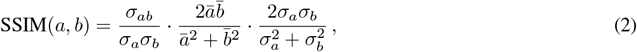

Where

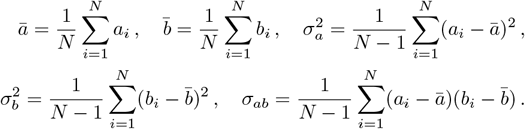

For conciseness, we refer to two data patches as *a* and *b*. For the case of comparing displacement fields, we evaluate SSIM loss as a combination of SSIM of the *x* and *y* displacement components, i.e.:

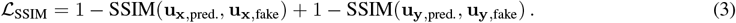

The first component of SSIM (Eq. (2)) is the correlation coefficient, the second measures the luminance similarity, and the third measures the contrast similarity. After being calculated for each patch, the SSIM scores are averaged across the image. The SSIM is normalised to [-1, 1], with higher scores indicating greater similarity between images. Therefore, for use as a loss function, the SSIM is subtracted from one. The SSIM implementation was based on previous work [44].

(3) LPIPS is emerging as a powerful loss that aligns with human estimates of image similarity [29]. To compute it, first, a small pre-trained AlexNet model [45] is evaluated upon corresponding patches of the input images. For each pixel within these patches, the feature embeddings produced by each layer of the network are normalised channel-wise and weighted using a learned vector. At each layer, the *l*_2_ distance between the resultant vectors originating from each patch is computed, and then averaged across all layers and pixels to give an overall patch score. The final metric is the mean score of all image patches. As with the SSIM, for use as a loss function, the LPIPS is subtracted from one. Here, an implementation of the LPIPS loss provided by the open-source `torchmetrics` package [46] was used.

(4)To compute the rewarp loss, the deformed frame (Def.) is warped to approximate the reference frame (Ref.) using the inverse of network’s displacement estimates (−**u**_pred._). This is achieved using the same method as that used for simulating the deformed image (Section 5.2). To mimic the correlation coefficient used in DIC as a loss function, we calculate the negative of the Pearson Correlation Coefficient (NPCC):

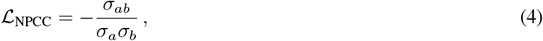

using the symbols defined in Eq. 2, where *a* is the real reference and *b* is the ‘rewarped’ deformed image.

### 5.4 DEFORM-Net software

We include the complete source code, implemented in PyTorch, as well as compiled versions of the network inference for open and rapid dissemination. The source code can be found at https://github.com/philipwijesinghe/displacement-estimation-for-microscopy. The compiled software, models and datasets can be found at https://doi.org/10.17630/feab7fa3-d77b-46e8-a487-7b47c760996a [47]. The original *Drosophila* data can be accessed in [26]. Details about the data and software are included in the Supplementary Information.

A major challenge in sharing deep learning methods is the requirement for particular development environments, hardware (specifically graphics processing unit (GPU)) and associated CUDA platforms. Training a new model without a GPU is unadvised due to exceptionally long training times. We provide open source displacement data simulation, PyTorch code and relevant environment specifications for those with the relevant hardware and those familiar with such methods.

However, inference using pre-trained models can be quick and accessible. Thus, we simplify the step of network inference in two ways. First, we provide a compiled version of the inference software and pre-trained models which can be executed on most consumer hardware using a command-line interface. Second, we provide an ImageJ/FIJI distribution which can perform this inference within its interface. Further documentation on the use of the software is provided in the Supplementary Information.

### 5.5 Optical flow

When considering a sequence of images, optical flow may be defined as the apparent motion of objects within a scene relative to the viewer. Optical flow results from the combination of objects’ actual motion and the motion of the camera. In many situations, accurately determining the optical flow allows objects’ true displacement between frames to be calculated. In the case of microscopy images, the camera is stationary, so determining the in-plane displacement of subjects only requires scaling the optical flow.

The brightness of pixels within an image may be represented by an intensity function *I*(**x**, *t*). The foundation of classical methods of determining optical flow (optical flow methods) is the brightness constancy assumption

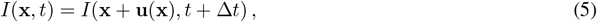

stating that the brightness of pixels representing features remains constant as those features move. **u**(**x**) is the displacement field determined by the projection of the three-dimensional motion of the scene into the image plane, and Δ*t* is the time-step between images. Applying a 1st order Taylor expansion leads to

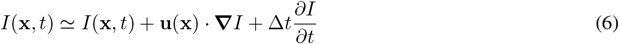

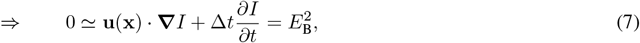

where 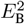 is commonly known as the brightness constraint. However, this equation is under-determined. Thus, to find the **u**(**x**) which minimises the brightness constraint additional assumptions must be introduced. Different optical flow methods are primarily distinguished by the additional assumptions they make.

The Lucas-Kanade algorithm [48, 49] splits the field into small subsets and requires that **u**(**x**) is homogeneous within them. The Horn-Schunck algorithm [50] is similar, but only stipulates that neighbouring pixels must have similar displacements. It adds a smoothness constraint, 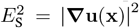, to Equation 7, and uses a variational approach to minimise 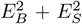 over the entire image. For our experiments, the LV-L1 method [33] was used. This is a improvement of the Horn-Schunck algorithm, modified to allow discontinuities in **u**(**x**) while maintaining the same principles.

These gradient-based classical optical flow methods are fast to compute, but the accuracy of the displacement estimates they produce is inherently limited. Their reliance on a truncated Taylor expansion means that the actual displacements must be sufficiently small for this approximation to be valid. Although image pyramids or alternative brightness constraint formulations [51] may be used to attempt to circumvent this limitation, this comes at the cost of computation time, and is not always wholly effective [16]. Moreover, the additional constraints introduced often relate the displacements of neighbouring pixels to one another, limiting the effective resolution of the results.

For this work, we used an implementation of the LV-L1 method [52] in an image pyramid form provided by `scikit-image` [53]. We found this resulted in the best performance metrics in supervised test data compared to other conventional methods, including the Lucas-Kanade and Horn-Schunck algorithms implemented natively in MATLAB (MathWorks, USA) and Farneback algorithm implemented using `opencv-python` [54].

### 5.6 Digital image correlation

Digital Image Correlation (DIC) is a related but distinct method of determining the displacement of objects within images between frames. The deformed image is split into patches, known as subsets. Each subset is then transformed in such a way as to maximise its correlation to the corresponding location on the reference image. This search process is iterative and slow. Typically, the process is sped up by determining an estimate for the translation of a chosen seed patch by computing its cross-correlation across the entirety of the reference image. This estimate is then iteratively refined, while allowing for affine deformation within the patch. Subsequent patches are then matched by progressively moving outwards from the seed. This significantly reduces the computational demand by computing the cross-correlation locally.

The accuracy of patch matching is dependent on the strength and distinctiveness of the texture within each patch; if repetitive patterns are present, false maxima in the cross-correlation can lead to incorrect displacement estimates. To counter this, speckle patterns are often applied to the objects, which can enable greater sub-pixel accuracy.

DIC offers several advantages over optical flow methods; for example, it is not inherently limited to small displacements, and the resolution of its estimates is flexible through the choice of the subset size and spacing (stride). Overlapping subsets can yield accurate, full-resolution estimates. However, in practice, it is more common to sacrifice resolution in order to speed up the process. Modern implementations, such as the open-source `ncorr2` [32], employ various optimisation strategies; nonetheless, processing is still substantially slower than gradient-based optical flow methods due to the iterative refinement performed for each patch in series.

We used `ncorr2` in this work. Specifically, we have adapted the MATLAB implementation [32] to automatically evaluate the `ncorr2` algorithm on image pairs. we used a spacing of 3 and subset size of 30, which we have experimentally found to produce, both, a reliable estimate in biological data and a high spatial resolution. DIC on a pair of images took approximately 120 s. Larger subsets resulted in lower spatial resolution, while smaller subets generated substantial artefacts. Several frame pairs from the 1,000+ frames generated artefacts and were automatically excluded from the evaluation of performance metrics.

### 5.7 Biological data

All biological videos were shared as part of previous studies or previously published work by others. No new biological experiments were carried out for this work.

Neonatal cardiomyocytes were obtained from postnatal day three wildtype mice as part of ongoing and previously published work in cell contractile sensing [55, 24, 25]. Briefly, the hearts were extracted from the pups and transferred to a dish containing ice-cold, calcium- and magnesium-free Dulbecco’s phosphate-buffered saline (DPBS) to preserve tissue integrity. The tissue was cut into pieces and placed in a falcon tube with papain (ten units per mL) and incubated for 30 minutes at 37 ^*°*^C with continuous rotation. Gentle reverse pipetting was utilised to disperse the tissue. A prewet cell strainer (70 *μ*m nylon mesh) was used to filter the cell suspension and remove large tissue debris. The cell suspension was centrifuged at 200*×*g for 5 minutes and resuspended into cell culture media (Dulbecco’s Modified Eagle’s Medium (DMEM) with 25 mM glucose and 2 mM Glutamax, 10% (v/v) FBS, 1% (v/v) non-essential amino acids, 1% (v/v) penicillin/streptomycin). Cells were pre-plated on uncoated petri dishes for 1–3h, this step removes fibroblasts and epithelial steps which will adhere to the uncoated dish. They were then counted and plated in a gelatin and fibronectin coated ibidi dish at a density of 6*×*10^4^ cells per cm^2^. Cell cultures were allowed to settle for 24h and kept in a humidified incubator at 37°C, with a 5% CO_2_ concentration. For widefield fluorescence imaging, cells were stained with 1000 nM SiR-actin (in DMEM with 10% FBS for 1h at 37°C), a dye that specifically binds to F-actin filaments in live cells, enabling real-time visualisation of the actin cytoskeleton. Cells were imaged 2 days post extraction. In the videos used in this work, 6–8 neonatal cardiomyocytes are visible in the field of view. The same cells were imaged sequentially using the different imaging modalities at an acquisition rate of 50 Hz.

The *Drosophila* LEC data were previously published in Pulido Companys et al. (2020) [26]. Briefly, LECs of the posterior compartment overexpress GMA-GFP, a fluorescent marker for F-actin [56], which enables visualisation of their pulsed contractions (genotype: *hh*.*Gal4>UAS*.*gma-GFP*). In the video used for this work, two cells are visible in the field of view. The time interval between frames is 10 s. The additional *Drosophila* LEC video that was not used in any network training comprised a separate sample that instead expressed a fluorescently labelled Myosin II regulatory light chain (Sqh::GFP) (genotype: *sqh[Ax3]; sqh::GPF; sqh::GFP*) [57]. In the video used for this work, one cell is visible in the field of view. The time interval between frames is 1 s.

## Acknowledgements

We acknowledge Kishan Dholakia for hosting the work and providing the computational resources to carry out neural network training and inference. We acknowledge Marcel Schubert for supervising the ongoing work with cardiomyocytes and Matthias König for isolating the cardiomyocytes from neonatal mice. PW acknowledges the support of the Research Fellowship from the Royal Commission for the Exhibition of 1851.

## Author contributions

PW conceived the project. SW, PW developed the methodology. SW, PW carried out the work and generated the results. SC provided the cardiomyocyte data and the related biological input. MB provided the *Drosophila* data and the related biological input. JK provided expertise in optical flow methods. All authors analysed the data. SW, PW drafted the manuscript with input from all authors. All authors edited and reviewed the manuscript. GB, PW supervised the work.

## Data Availability

The source code can be found at https://github.com/philipwijesinghe/displacement-estimation-for-microscopy. The compiled software, models and datasets can be found at https://doi.org/10.17630/feab7fa3-d77b-46e8-a487-7b47c760996a [47]. The original *Drosophila* data can be accessed in [26]. See Methods and Supplementary Information for additional details. Any additional training data not included here (due to large file sizes) can be provided upon request.

