## Supplementary Information for "Estimating full-field displacement in biological images using deep learning"

#### Contents

**S1 Rewarp loss as an unsupervised metric** **2**

**S2 The role of supervised and unsupervised training loss functions** **3**

**S3 Generalisation of DEFORM-Net on open-source data** **4**

**S4 Software documentation** **6**

#### S1 Rewarp loss as an unsupervised metric

Naively, we can consider the unsupervised 'rewarp' loss as an error metric for real data. When a deformed image is 'undeformed' using an accurate estimate of displacement, it should correlate well to the reference image. This process could be thought of as similar to DIC. Figure S1(a), however, illustrates a major challenge in using such correlation as a metric of accurate displacement. Consider the displacement of a set of points between two frames. It is reasonable that for a small enough subset, there may be several possible displacements that lead to a high local correlation in intensities. Only when the subset size is sufficiently large to describe a unique set of image features, will the estimate likely possess a global maximum that corresponds to the true displacement field.

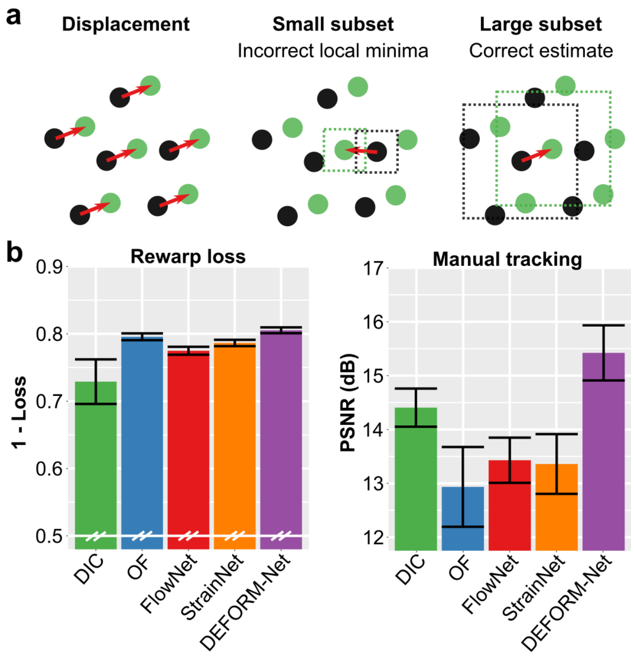

Figure S1: Rewarp loss as an error metric. (a) The challenge of using image intensity correlation as a metric for displacement. (b) Performance evaluated using the intensity correlation-based rewarp metric (n = 50) compared to manually tracked (n = 9) displacements. Error bars are standard error of mean.

Evaluating real experimental image frames of the *Drosophila* LECs dataset (F-actin) (n=50) using the ground-truth-free rewarp metric showed that, unexpectedly, DIC performs overall the worst (Fig. S1(b)). The scenario of evaluating rewarp loss on a pixel-by-pixel basis leads to the following property: an accurate displacement estimate improves rewarp loss, however, a good rewarp loss does not indicate an accurate displacement estimate. This idea of some constancy in intensity values during displacement is common to DIC and OF, and their solutions differ in the method of regularisation and constraints used to converge to a globally accurate solution. Thus, it is unsurprising that all methods intrinsically perform well in this loss, despite producing different estimates. This loss may be improved by evaluating correlations between local subsets instead of pixel by pixel. However, this pursuit will likely lead to solutions that are very similar to that used in DIC, with the associated challenges in computational speed.

#### S2 The role of supervised and unsupervised training loss functions

DEFORM-Net comprises several training loss functions to enhance its performance and generality. Here, we describe the selection of the losses and their utility with respect to performance metrics in simulated and real data. Table S1 lists the performance metrics of each loss in models with various combinations of training loss functions. We compare a baseline model, which utilised solely the conventional MSE and SSIM losses, against those with added rewarp loss or LPIPS loss, as well as the combined model with all losses included. These are also benchmarked against the conventional DIC and OF methods. The performance is evaluated using the PSNR, SSIM, LPIPS and rewarp metrics in simulated testing data, as well as in real *Drosophila* (F-actin) image frames compared to manual tracking (Manual PSNR). We also evaluate the performance of each model when all *Drosophila* images are specifically excluded from training (Cross-domain PSNR) to test generalisation.

Table S1: The role of training loss functions on performance evaluated on both simulated and real displacement data. Highest performance values within standard error bounds are highlighted in bold.

| **Method** | **PSNR (dB)** | **SSIM (/100)** | **LPIPS (/100)** | **Rewarp (/100)** | **Manual PSNR (dB)** | **Cross-domain PSNR (dB)** |
|---|---|---|---|---|---|---|
| DIC | 24.6 (0.2) | 68.5 (0.6) | 92.6 (0.4) | 97.1 (0.3) | 14.4 (0.4) | 14.4 (0.4) |
| OF | 19.9 (0.3) | 34.8 (1.4) | 87.1 (0.7) | 95.6 (0.5) | 13.0 (0.7) | 13.0 (0.7) |
| Baseline | 24.4 (0.2) | **77.4 (0.4)** | 95.4 (0.3) | 96.8 (0.3) | 14.7 (0.6) | **14.7 (0.6)** |
| Rewarp loss | **26.3 (0.2)** | 73.3 (0.5) | 95.8 (0.3) | **97.9 (0.2)** | **15.0 (0.5)** | **14.8 (0.6)** |
| LPIPS loss | 24.9 (0.3) | 75.2 (0.5) | **96.3 (0.2)** | 97.0 (0.3) | 14.5 (0.5) | 14.4 (0.5) |
| Combined loss | **26.5 (0.2)** | 73.7 (0.5) | **96.4 (0.2)** | **98.0 (0.2)** | **15.4 (0.5)** | **15.0 (0.5)** |

The baseline model performs adequately in all metrics and matches, for most part, the performance of DIC. Interestingly, with the addition of rewarp loss, the PSNR error in simulated testing data is improved substantially. Naively, we may assume that training with MSE loss should maximise PSNR ($\propto 1/\text{MSE}$). However, because rewarp loss is unsupervised and describes an apriori physical property of the data, it likely helps the network avoid overfitting, and thus, improves performance when evaluated in unseen testing data. Despite this, this benefit is only conferred marginally to the performance in real data. Alternatively, the addition of LPIPS loss to the baseline model has little impact on the network performance.

Interestingly, when both rewarp and LPIPS losses are included in a combined model, the performance is substantially optimised in both simulated and real data across most metrics. Based on the previous performance of the rewarp metric compared to manual tracking (Supplementary Note S1), we can see that the rewarp loss can be challenged by the presence of local minima. This will likely result is substantial high-spatial-frequency local displacements in the estimate that would not be present in the simulated ground truth data. The added LPIPS loss likely focuses its attention on these details, while MSE or even SSIM losses are known to not be sensitive to such high-frequency content [1]. While the baseline model is adequate at meeting the DIC challenge, we have utilised the combined model for all other demonstrations in this work.

#### S3 Generalisation of DEFORM-Net on open-source data

Here, we demonstrate the performance of DEFORM-Net on a publicly available dataset with attention to the role of microscopy-informed noise in model training. Lye et al. [2] provide open-source (CC-BY-4.0) multimodal confocal microscopy data [3] of a *Drosophila* embryo undergoing gastrulation, followed by the beginning of germband extension. Figures S2(a–c) show a representative frame of the two fluorescence channels and the composite image of Myosin II (sqh-GFP[KI]) and cell membranes (Gap43-mCherry), respectively. Gastrulation and germband extension are dynamic, multi-cell processes during embryonic development. During gastrulation, the mesoderm cells invaginate, forming the ventral furrow. This is followed by germband extension, where ectoderm cells converge and extend along the anterior-posterior axis (left-to-right).

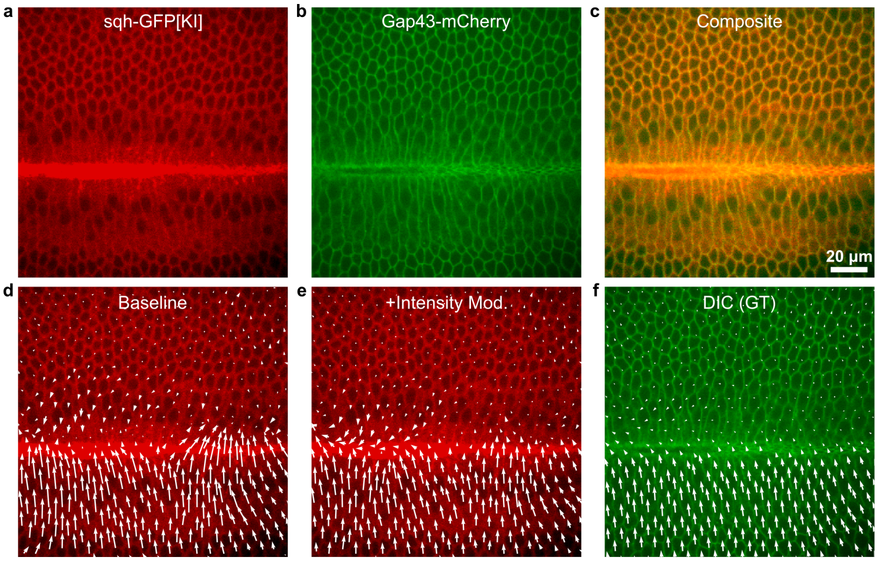

Figure S2: Performance and generalisation of DEFORM-Net on publicly available multimodal microscopy data provided in Lye et al. [2, 3]. A frame from a video of germband extension of a *Drosophila* embryo undergoing gastrulation, followed by the beginning of germband extension. Cells express (a) Myosin II-GFP (sqh-GFP[KI]) and (b) Gap43-mCherry (a cell membrane marker), and (c) a composite of the two channels. (d, e) Vector plots evaluated on the poorer signal Myosin II channel in (a) using (d) a baseline network with MSE and SSIM losses and (e) the same network with added intensity modulation during training; compared to a ground truth estimated using DIC on the higher signal cell membrane contrast in (b).

Table S2: Performance of DEFORM-Net on publicly available multimodal microscopy data provided in Lye et al. [2]. Peak signal-to-noise ratio (PSNR) of the MSE is provided with respect to the DIC estimate as well as compared to manual tracking ($n = 4$). Highest performance values within standard error bounds are highlighted in bold.

| **Method** | **PSNR (vs. DIC) (dB)** | **Manual PSNR (dB)** |
|---|---|---|
| DIC | - | **16.0 (1.0)** |
| OF | 19.0 (0.3) | **15.3 (1.0)** |
| Baseline | 18.0 (0.1) | 14.0 (0.7) |
| Baseline + Int. Mod. | 20.0 (0.1) | **15.6 (1.0)** |
| DEFORM-Net | **20.6 (0.1)** | **15.5 (0.9)** |

The data provided in Lye et al. [2] is particularly useful for several reasons. First, the data features distinct full-field displacement of tissue with good feature contrast and with regular timelapse sampling of 30 s per frame (140 frames). Second, the registered and simultaneously acquired fluorescence channels comprise a relatively poor SNR and contrast data of Myosin II, which we use to evaluate DEFORM-Net, and a higher SNR and contrast data of the cell membranes, which we use to generate estimates of the ground truth displacement. Specifically, we perform DIC on the cell membrane data and assume that this is a close estimate of the ground truth displacement (Fig. S2(f)). We then evaluate several DEFORM-Net networks in their ability to estimate similar motion from the poorer contrast Myosin II data. Specifically, we evaluate a 'baseline' network trained solely on cardiomyocyte data and the MSE and SSIM losses (Fig. S2(d)), a ‘baseline’ network with the addition of the ‘intensity modulation’ noise term (Fig. S2(e)), and the DEFORM-Net network that comprises all data, noise models, and losses, as reported in the main manuscript. By inspecting Figs S2(d) and (e), we see that the network without ‘intensity modulation’ struggles in estimating accurate displacement vectors in the vicinity of the bright horizontal feature, which is the folding of the cells out of the plane of focus.

Table S2 quantifies the errors between these networks; first, by taking the MSE between the estimate and the DIC estimate from the cell membrane channel, presented as the PSNR (vs. DIC); second, by measuring the PSNR of MSE using manual tracking ($n = 4$), as was done in the main manuscript. We can see that for both error estimates, the addition of ‘intensity modulation’ in training resulted in significantly improved performance in unseen data. The final DEFORM-Net network comprising both cardiomyocyte and *Drosophila* LEC data, and all losses, resulted in closer correspondence to DIC estimates, while not offering significant differences in performance when compared to manual tracking. In all cases, addition of ‘intensity modulation’ was valuable in this particular biological data.

### S4 Software documentation

DEFORM-Net can be interacted with in several ways. We release the code as open source at:
`https://github.com/philipwijesinghe/displacement-estimation-for-microscopy`

The associated data, compiled code and trained models can be found in our institutional repository:
`https://doi.org/10.17630/feab7fa3-d77b-46e8-a487-7b47c760996a` [4].

DEFORM-Net is implemented in Python using the PyTorch framework. Because training new models requires both a familiarity with Python-based deep learning and potentially expensive hardware, we also release our pre-trained models and several options for inference (processing of new data with existing models). This includes inference using the deepImageJ plugin [5] for the widely used ImageJ/FIJI software [6], as well as a command-line interface for using standalone compiled binary files (Windows).

Importantly, inference requires a pre-trained model. The pre-trained models that we make available as part of this work are described in Section S4.1. Further, inference using compiled binaries or training and inference using python require the input data to be standardised in a particular format and folder layout, which is described in Section S4.2. Minimal example data and the biological data used in this paper are described in Section S4.3.

#### S4.1 Pre-trained models

Pre-trained models can be found on our institutional repository [4]. Each model is released in two formats: (1) as a PyTorch state dictionary (which can be loaded by our Inference Class in python), and (2) as TorchScript models (which can be loaded by deepImageJ).

We release the following models:

- `deformnet`: model that combines all loss functions and biological training data;
- `deformnet-base`: model that comprises only the MSE and SSIM losses for all training data;
- `deformnet-cardio`: model that was trained on cardiomyocytes only;
- `deformnet-droso`: model that was trained on drosophila only.

Due to file upload limits, we provide the individual `deformnet` model in pytorch and torchscript format (`model-deformnet-pytorch.zip` and `model-deformnet-torchscript.zip`). All four models are further provided in a combined zipped format for pytorch and torchscript (`models-pytorch.zip` and `models-torchscript.zip`).

#### S4.2 Common data structure

Programmatic interaction with DEFORM-Net requires the data to be in a particular file and folder structure. Inference requires a folder where the reference and deformed image pairs are placed in respective subfolders called 'Ref' and 'Def'. This may be a single image pair, multiple image pairs, or an image sequences reconfigured as sequential image pairs. Image pairs do not have to share a common name, but should be ordered. Video sequences can be saved as an image sequence (a sequence of individual images) in ImageJ and image indices 1:N-1 may be saved as Ref and 2:N may be saved as Def. For example:

```
/drosophila/Ref/img_0001.tif
               /img_0002.tif
           /Def/img_0002.tif
               /img_0003.tif
```

will pair `img_0001` with `img_0002`, and `img_0002` with `img_0003`.

Training requires similar 'Ref' and 'Def' folders as well as 'Dispx' and 'Dispy' folders with corresponding $x$ and $y$ displacement matrices in numpy (.npy) format. These can be generated using our displacement simulation code (Section S4.6). Further, an appropriate partition of training and validation data must be provided (testing data is optional). For example:

```
/training/drosophila/training/Ref/
                             /Def/
                             /Dispx/
```

```
                /Dispy/
    /testing/ "
    /validation/ "
```

Inference will generate a new pair of folders in the parent directory, called ‘Dispx-inference’ and ‘Dispy-inference’, which will contain numpy matrices of estimated $x$ and $y$ displacement fields for each image pair. An example of folder structures can be found in our minimal example data (Section S4.3).

##### S4.3 Data

Datasets used in this work are provided on our institutional repository [4]. Specifically, we provide a minimal example dataset for inference and training, demonstrating the typical data layout, configurations files and outputs (`minimal_example_data.zip`). Further, we provide the cardiomyocytes videos as image sequences (`data-cardiomyocytes.zip`). The *Drosophila* videos may be accessed from the original publication [7].

We further provide a portable snapshop of Fiji (`Fiji.app.zip`) that includes deepImageJ and our models. We also provide the standalone compiled binaries for inference (`deformnet_inference_cli.zip`).

We do not provide all of the simulated training data due to size (10–100 GBs), however, it is possible to regenerate the data from the raw biological videos and the displacement simulation code provided. Further, specific training data may be shared upon reasonable request.

##### S4.4 Inference using deepImageJ

DeepImageJ is an ImageJ/FIJI plugin that enables inference using compatible TorchScript models, such as those available from the BioImage Model Zoo [5]. Installation of deepImageJ can be found on `https://deepimagej.github.io/index.html`. Our TorchScript DEFORM-Net models (provided as .zip files) may be directly installed by the plugin (via Plugins > DeepImageJ > DeepImageJ Install Model, as a Private Model).

DEFORM-Net may be then run via Plugins > DeepImageJ > DeepImageJ Run. However, it requires the input image to be of size (channels = 6, height, width), where the channels comprise the RGB components of the reference image followed by the RGB components of the deformed image. Further, the data must be a 32-bit float normalised to [0, 1].

Note: Our example models in torchscript are formatted for image inputs of $1024 \times 1024$. While deepImageJ can support tiling and stitching, we note that there are unintended artefacts in the present implementation. Thus, we recommend to format the input image stack to have an image size of $1024 \times 1024$, or factors thereof. Smaller images may be padded with zeros using the ImageJ ‘Adjust > Canvas Size’ tool.

Because the input shape is not a conventional format, we also provide an ImageJ macro that will sequentially process an image series (8-bit grayscale images) or video and output corresponding $x$ and $y$ displacement fields.

This macro is listed below and is available on our github repository. The macro may be edited to select the appropriate model name, image normalisation, etc.

```
// Start with a video stack in imagej
model="deformnet"  // SELECT MODEL NAME AS INSTALLED IN DEEPIMAGEJ <!>
name=getTitle();
getDimensions(w, h, channels, slices, frames);

for (i = 1; i < slices; i++) {
        selectImage(name);
        i2 = i + 1;

        // Create network input
        run("Make Substack...", "slices="+i+","+i+","+i+","+i2+","+i2+","+i2);
        rename("img_pair_"+i);

        // normalise to 0,1 (Assumes 8-bit Grayscale; EDIT THIS SECTION FOR YOUR DATA
            TYPE) <!>
        run("32-bit");
        run("Divide...", "value=256 stack");
        setMinAndMax(0, 1);
        run("Properties...", "channels=6 slices=1 frames=1 pixel_width=1.0000
            pixel_height=1.0000 voxel_depth=1.0000");
```

```
        // Run model
        run("DeepImageJ Run", "model="+model+" format=Pytorch preprocessing=[no
            preprocessing] postprocessing=[no postprocessing] axes=C,Y,X tile
            =6,1024,1024 logging=Normal");

        // Format output and append to stacks
        rename("output_"+i);
        run("Split Channels");

        selectImage("C1-output_"+i);
        rename("ux_output_"+i);
        run("Grays");
        if (i==1) {
                rename("ux_output");
        } else {
                run("Concatenate...", "open image1=ux_output image2=ux_output_"+i+"
                    image3=[-- None --]");
                rename("ux_output");
        }

        selectImage("C2-output_"+i);
        rename("uy_output_"+i);
        run("Grays");
        if (i==1) {
                rename("uy_output");
        } else {
                run("Concatenate...", "open image1=uy_output image2=uy_output_"+i+"
                    image3=[-- None --]");
                rename("uy_output");
        }

        // Cleanup
        selectImage("img_pair_"+i);
        close();
}
```

##### S4.5 Inference using compiled binaries

We have compiled our inference code using ‘pyinstaller’. The complied binaries feature the required environment for inference, including the relevant CUDA environments and python packages, such that it may be run standalone on a Windows system. The files are available from our institutional repository [4] (`deformnet_inference_cli.zip`). The main point of entry is the `deformnet_inference.exe` executable. This may be run using the command line with identical arguments to `deformnet_inference.py` (Section S4.6).

At the minimum, one must specify the location of the PyTorch model state dictionary and the parent folder, as described in Section S4.2. For instance:

```
deformnet_inference.exe "G:/example/models/example_model/saved_model/
    checkpoint.pt" "G:/example/inference/drosophila" --crop 1024
```

##### S4.6 Training and inference using PyTorch

Training and inference using PyTorch requires an appropriate Python environment and training data or pre-trained models.

###### S4.6.1 Environment

DEFORM-Net was tested using PyTorch 2.2.1 (CUDA 12.1 (NVIDIA driver >=528.33); Python 3.11) managed using miniconda (or Anaconda) package manager. Detailed installation requirements are listed in ENVIRONMENT.md.

##### S4.6.2 Generating training data

Training data in the format required by DEFORM-Net (Section S4.2) can be generated from a series of images using the fractal Perlin generation method. An example script is provided in `deformnet_prepare_data.py`. Specifically, the `in_path` must point to a folder containing a series of microscopy images, which may be of various sizes or from multiple sources.

##### S4.6.3 Training

An example script for training DEFORM-Net is provided in `deformnet_train_model.py`. Training relies on a user-editable configuration file (`config.yml`), which may be edited with any text editor. The user must place this configuration file in a desired folder where the model will later be stored. An example configuration file is provided in the sample dataset (Section S4.3). The user must then edit the python code to specify the directory of the configuration file.

##### S4.6.4 Inference

Example scripts for DEFORM-Net inference are provided in a script format `deformnet_inference_script.py` or in a command line interface format `deformnet_inference.py`.

```
    usage: deformnet_inference.py [-h] [--suffix SUFFIX] [--crop CROP] [--
        overwrite OVERWRITE] [--save_fmt SAVE_FMT]
                                 model data

positional arguments:
  model                 full path to trained model
  data                  full path to paired data for inference

options:
  -h, --help            show this help message and exit
  --suffix SUFFIX       suffix to append to inference outputs
  --crop CROP           crop data
  --overwrite OVERWRITE
                        overwrite output if it exists
  --save_fmt SAVE_FMT   save format: 'npy', 'tiff', or 'both'
```
